# Mutation spectra across 3,034 *Saccharomyces cerevisiae* genomes distinguish the Georgian Wine population

**DOI:** 10.64898/2026.09.28.755217

**Authors:** Cian Wheeler, Pengyao Jiang

**Affiliations:** Center for Mechanisms of Evolution, Biodesign Institute, School of Life Sciences, Arizona State University

## Abstract

Mutation spectra vary among populations, and a comprehensive view of this variation requires broad sampling and fine-scale population resolution. Here, we profile mutation spectra from single-nucleotide polymorphisms and small insertions and deletions in a uniformly processed collection of 3,034 globally sampled *Saccharomyces cerevisiae* genomes, and an extended dataset including the previously sampled global 1,011-genome collection. Major mutation spectrum patterns reported from the 1,011 genomes remain evident in the extended dataset, including the pronounced mutation-spectrum differentiation of African Beer strains. At finer population resolution, we identify additional variation among wine-associated populations, including a Georgian Wine-associated signal that is strongest among rare variants and is therefore consistent with a comparatively recent change in mutation processes. We further apply a sequence-context-aware classification to characterize small-indel profiles across the 3,034 natural yeast genomes. Together, our analyses show that expanded sampling and refined population assignments provide a more comprehensive view of mutation-spectrum variation within *S. cerevisiae*.

## Introduction

Mutations are the ultimate source of genetic variation, providing the raw material on which natural selection and genetic drift act (Wright 1931, 1930; Kimura 1983). Mutations arise through diverse molecular mechanisms, and biases in DNA replication, damage, and repair generate characteristic patterns of nucleotide substitutions and small insertions and deletions (Volkova et al. 2020). The relative frequencies of these changes and their sequence contexts constitute a mutation spectrum (Liu and Zhang 2021; Melde et al. 2022; Harris 2015; Hodgkinson and Eyre-Walker 2011). Since mutation spectra reflect the processes that generate and repair DNA lesions, variation in these spectra can provide insight into the molecular mechanisms shaping genome evolution (Carlson et al. 2020; Seplyarskiy et al. 2021).

Population genomic data retain a record of historical mutational input through variants segregating in natural populations. Analyses of mutation spectrum patterns from polymorphisms have revealed recent spectrum changes in humans, spectrum differentiation in mammal species, strain-specific differences in mice, and close agreement between polymorphism- and experiment-based spectra in yeast (Harris 2015; Harris and Pritchard 2017; Dumont 2019; DeWitt et al. 2021; Zhu et al. 2017; Goldberg and Harris 2022; Beichman et al. 2023). A complementary framework in cancer genomics decomposes observed mutation spectra into recurrent mutational signatures, allowing estimation of the contributions of multiple underlying mutational processes (Alexandrov et al. 2013, 2020). The Catalogue of Somatic Mutations in Cancer (COSMIC) provides standardized reference signatures, some of which have been associated with endogenous processes, exogenous exposures, or defects in DNA maintenance (Alexandrov et al. 2020).

The global diversity of *Saccharomyces cerevisiae* provides an ideal system for studying variation with genetically diverse isolates from wild and human-associated environments. The 1,011-genome collection defined major populations across the species (Peter et al. 2018). Analyses of that collection identified reproducible population- and strain-specific differences in substitution spectra. African Beer strains formed a pronounced outlier, while rare C>A-enriched variants distinguished individual strains (Jiang et al. 2021). Direct mutation measurements and genetic complementation linked the C>A enrichment in two Mosaic Beer strains to a natural mutator allele of *OGG1* (Jiang et al. 2021). Subsequent work found genomic patterns consistent with the long-term maintenance of mild-effect mutator alleles in African Beer and their transmission into a subset of French Dairy strains (Jiang et al. 2025). These studies demonstrated that extensive mutation spectrum variation can exist in natural isolates of *S. cerevisiae*.

Recently, the compilation of sequencing data from 3,034 *S. cerevisiae* isolates substantially expands both sampling and population resolution (Loegler et al. 2024). This resource integrates isolates from many studies under a common variant-calling pipeline, resolves the wine superclade into several subpopulations, and provides both single-nucleotide and small-indel calls. This dataset allows us to test whether mutational patterns identified in the 1,011 genomes persist in a larger dataset and whether novel mutational signatures can be discovered in this extended dataset.

Small insertions and deletions provide a complementary view to single nucleotide mutations. In humans, population-scale sequencing and well-developed classification frameworks have linked indel variation to repeat structure, local sequence features, and selective constraint (Montgomery et al. 2013; Koh et al. 2025). In yeast, polymerase slippage and mismatch repair are established determinants of short-indel formation (Harfe and Jinks-Robertson 2000; Gragg et al. 2002), and rare polymorphisms have been used to examine indel context and selective filtering (Zhu et al. 2017). However, direct estimates of spontaneous indels in *S. cerevisiae* have come mainly from mutation-accumulation studies (Zhu et al. 2014; Sharp et al. 2018), and systematic, sequence-context-resolved comparisons among natural yeast populations remain limited.

Here, we analyze mutation spectra from the 3,034-genome resource of natural polymorphisms and small indels. The expanded dataset retains the major patterns from the previous analysis with the 1011 genomes while revealing additional differentiation among wine-associated populations, including a Georgian Wine signal that is strongest among rare variants. We also adapt a sequence-context-aware classification of small indels to test whether population differentiation extends beyond single-nucleotide substitutions. Together, these analyses revealed additional mutation-spectrum variation from standing genetic variation in *S. cerevisiae* from this extended dataset.

## Results

### Mutation signature decomposition from the 3034 *S. cerevisiae* genomes reveals a novel signature among wine clades

We analyzed mutations from polymorphisms in the 3034 *S. cerevisiae* natural isolates (Loegler et al. 2024), the largest genomic collection of *S. cerevisiae* to date. Similar to the analysis with 1011 *S. cerevisiae* (Jiang et al. 2021), we included SNPs polarized by the same *S. paradoxus* as the outgroup, and excluded closely related strains using the same criteria, downsampling to no more than 30 strains per group, leaving a final set of 785 non-closely related strains after excluding strains with extensive introgression from *S. paradoxus* (Figure 1A, Materials and Methods). We counted single-nucleotide substitution mutations as well as triplet mutations that include neighboring sites. To discern any unique mutation signature within *S. cerevisiae*, we performed mutation signature decomposition using SigProfilerExtractor on the triplet mutations (Islam et al. 2022).

**Figure 1:**
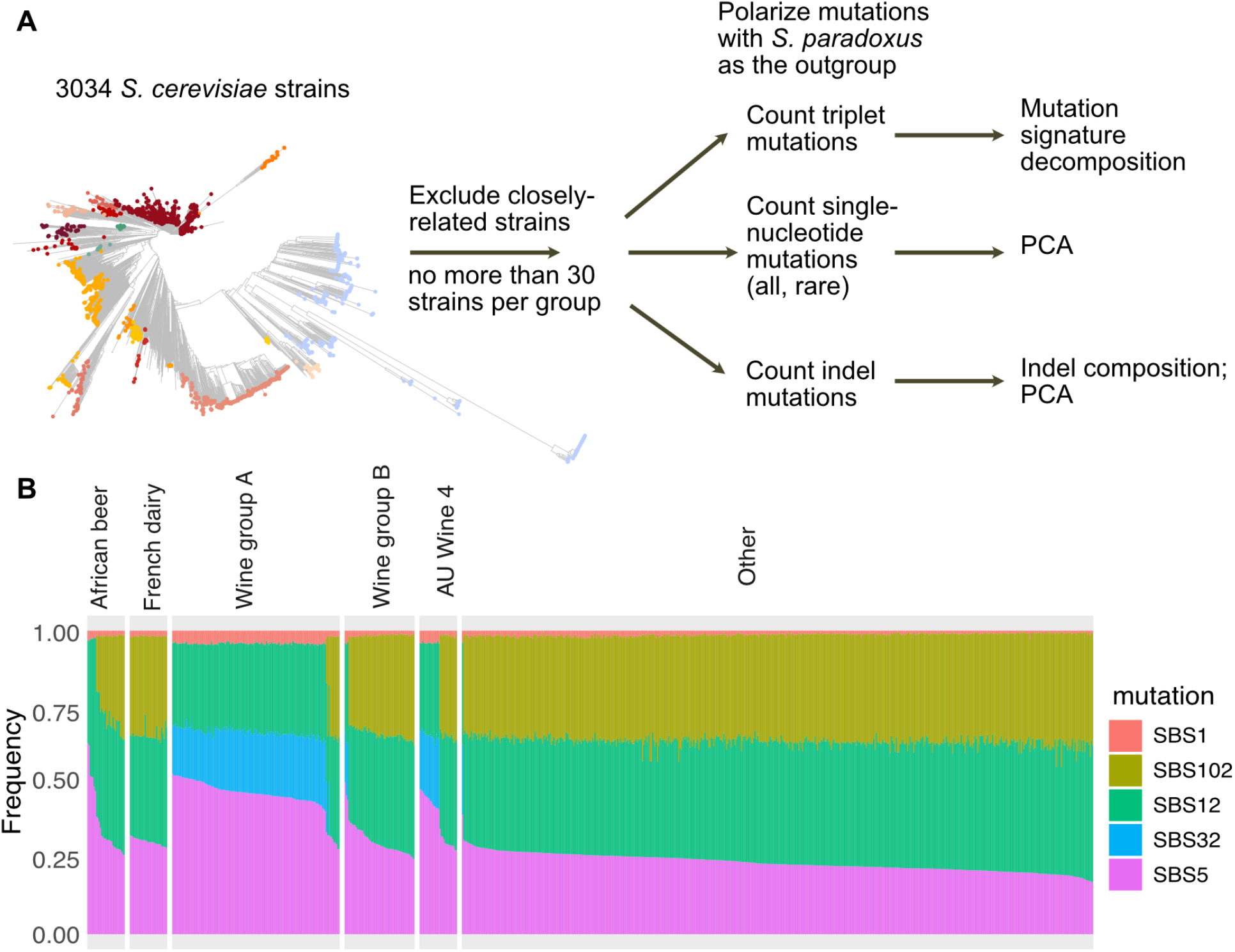
*S. cerevisiae* signature decomposition reveals a novel signature among wine clades: (A) Analysis flow in this work. Natural polymorphisms and indels were extracted from the 3034 *S. cerevisiae* dataset from non-closely related strains and were further subjected to mutation analysis. (B) Mutation signature decomposition of triplet mutation spectrum via SigProfiler. Using known COSMIC signatures, a breakdown of contributions was predicted based on the 96-triplet mutation context. Wine group A (AU Wine 1-3, Georgian Wine, and West Europe Wine) exhibits different signatures than Wine group B (AU Wine 5-6, and Commercial Wine), while AU Wine 4 was split more equally between signature profiles.

We identified five major COSMIC signatures across the 3034 *S. cerevisiae* natural isolates — SBS1, SBS102, SBS12, SBS32, and SBS65 (Figure 1B). Consistent with a recent analysis of the 1011 *S. cerevisiae* that a subset of African Beer strains lack the SBS102 signature but have an increased proportion of SBS5 (Jiang et al. 2025), we recapitulate the same structure in this extended dataset (Figure 1B). In addition, we discovered a novel signature in the wine superclade that was not present in the 1011 *S. cerevisiae* genomes (Jiang et al. 2025): the SBS32 signature replacing SBS102 was observed (Figure 1, Supplementary Figure 1). With the inclusion of more samples in the 3,034-genome collection, the wine superclade was further divided into subgroups (Loegler et al. 2024), rather than treated as a single large group as in the 1,011-genome

*S. cerevisiae* collection (Peter et al. 2018). SBS32 predominantly consists of C>T mutations (75%), in combination with C>A, A>T, and A>G mutations (Inman et al. 2018). Based on the presence or absence of SBS32, we split the wine superclade into two main groups: Wine group A and Wine group B. Wine group A consists of AU Wine 1-3, Georgian Wine, and West Europe Wine, which are characterized by the additional SBS32 signature and the absence of SBS102. Not all strains within each subgroup exhibit SBS32, but the majority do (Supplementary Figure 1). Wine group B consists of AU Wine 5–6 and Commercial Wine; most strains in these subgroups lack SBS32. AU Wine 4 is an exception, which cannot be placed in either group, since roughly half of the strains in that group have the SBS32 signature, while the other half do not. For simplicity of reporting results, we will use the new annotation of Wine group A and Wine group B unless particular subgroups stand out. However, the analyses themselves were performed using the original group annotations from (Loegler et al. 2024).

### The mutation spectrum from 3034 *S. cerevisiae* genomes shows strong similarity to the 1011 *S. cerevisiae* strains

We performed mutational Principal component analysis (PCA) on the 3034 *S. cerevisiae* natural isolates to further discern mutation-spectrum patterns in this extended dataset. The mutational PCA was performed similarly to that for the 1,011-genome collection (Jiang et al. 2021), except that we included strains with extensive introgression from *S. paradoxus* for completeness in this analysis, with a total of 858 strains (Peter et al. 2018) (Materials and Methods). Initial analysis of the mutation spectra showed that the PCA is strongly skewed by A>G mutations in the African Beer population, with PC1 driven primarily by the contrast between A>G and all other mutation types (Supplementary Figure 2). The observation that African Beer is an outlier is consistent with previous studies of the 1,011 *S. cerevisiae* collection (Jiang et al. 2021) and other *Saccharomyces* species (Jiang et al. 2025). This skew highlights the substantial difference in mutation spectra between African Beer and the other populations. Because A>G is one of the more frequent mutation types, it receives greater weight in the analysis, which could distort the PCA as more African Beer strains are included. To characterize variation equally among higher- and lower-frequency mutation types, we therefore repeated the PCA after normalizing each mutation type (Materials and Methods).

**Figure 2:**
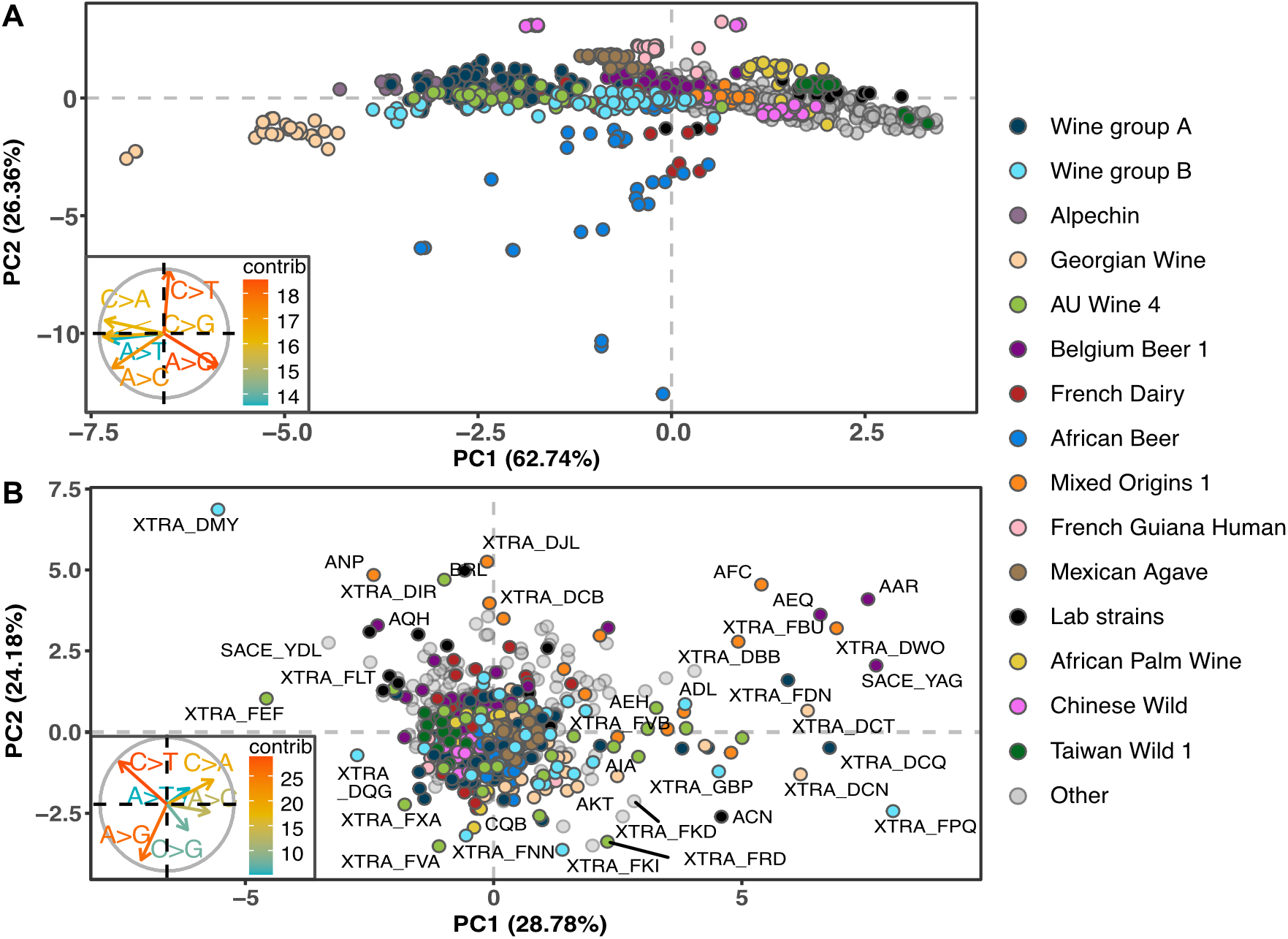
Mutation spectra of 3034 *S. cerevisiae* isolates uncover population- and strain-specific patterns: (A) Mutational PCA of *S. cerevisiae* variants from all polarized SNPs with<50% allele frequency. Each point represents a single strain, with color used to define their clades. African Beer, Chinese Wild, and Georgian Wine are outliers. (B) Mutational PCA of rare variants in the dataset.

After normalization, the mutational PCA recapitulated an overall structure similar to that found previously using the 1,011 *S. cerevisiae* genomes, with the main difference that PC1 and PC2 were interchanged (Figure 2A, (Jiang et al. 2021)). PC1 now formed a gradient along which most strains were distributed, whereas PC2 separated the outlier groups, including African Beer. African Beer strains were enriched for A>G and A>C mutations, whereas French Dairy strains formed two clusters. One clustered near the other *S. cerevisiae* strains, while the other exhibited an intermediate mutation spectrum between African Beer and other *S. cerevisiae* strains (Jiang et al. 2025). At the opposite end of PC2 from African Beer, Chinese Wild strains were enriched for C>T transitions (Figure 2A). We observed a similar mutation-spectrum pattern in the Taiwanese population in the 1,011-strain collection. Upon further inspection, however, we found that this similarity was simply due to a different grouping scheme and naming convention. In the 3,034-genome dataset, (Loegler et al. 2024) grouped the strains originally classified as Taiwanese in the 1,011-genome collection with Chinese Wild rather than Taiwan Wild 1. We also observed the Georgian Wine clade as an outlier along PC1, a pattern that has not been reported previously.

We performed additional PCA analysis with various modifications to the subsampling and grouping criteria and mutation types, and obtained largely consistent patterns throughout. To ensure subsampling of strains of each group did not affect overall patterns, we also performed mutational PCA of all the non-closely related strains without subsampling strains from each group and obtained similar results (Supplementary Figure 3). The relative positions of strain-specific mutation profiles did not change when we repeated mutational PCA after excluding the introgressed strains (Supplementary Figure 4). We also performed mutational PCA on triplet mutation types (Supplementary Figure 5). The resulting pattern is largely consistent with that based on single-nucleotide mutation types, except that Georgian Wine was shifted further toward negative PC2 values. This discrepancy was more pronounced in the PCA performed without subsampling strains from this group (Supplementary Figure 6). We suspect that this may be due partly to the normalization procedure for triplet mutations because not every mutation category had sufficient counts. Mutation types with lower counts may be more susceptible to artifacts introduced during normalization. We therefore performed an additional analysis without normalizing the triplet mutation counts, and the skew of Georgian Wine toward negative PC2 values was no longer observed (Supplementary Figure 7), supporting our hypothesis.

**Figure 3:**
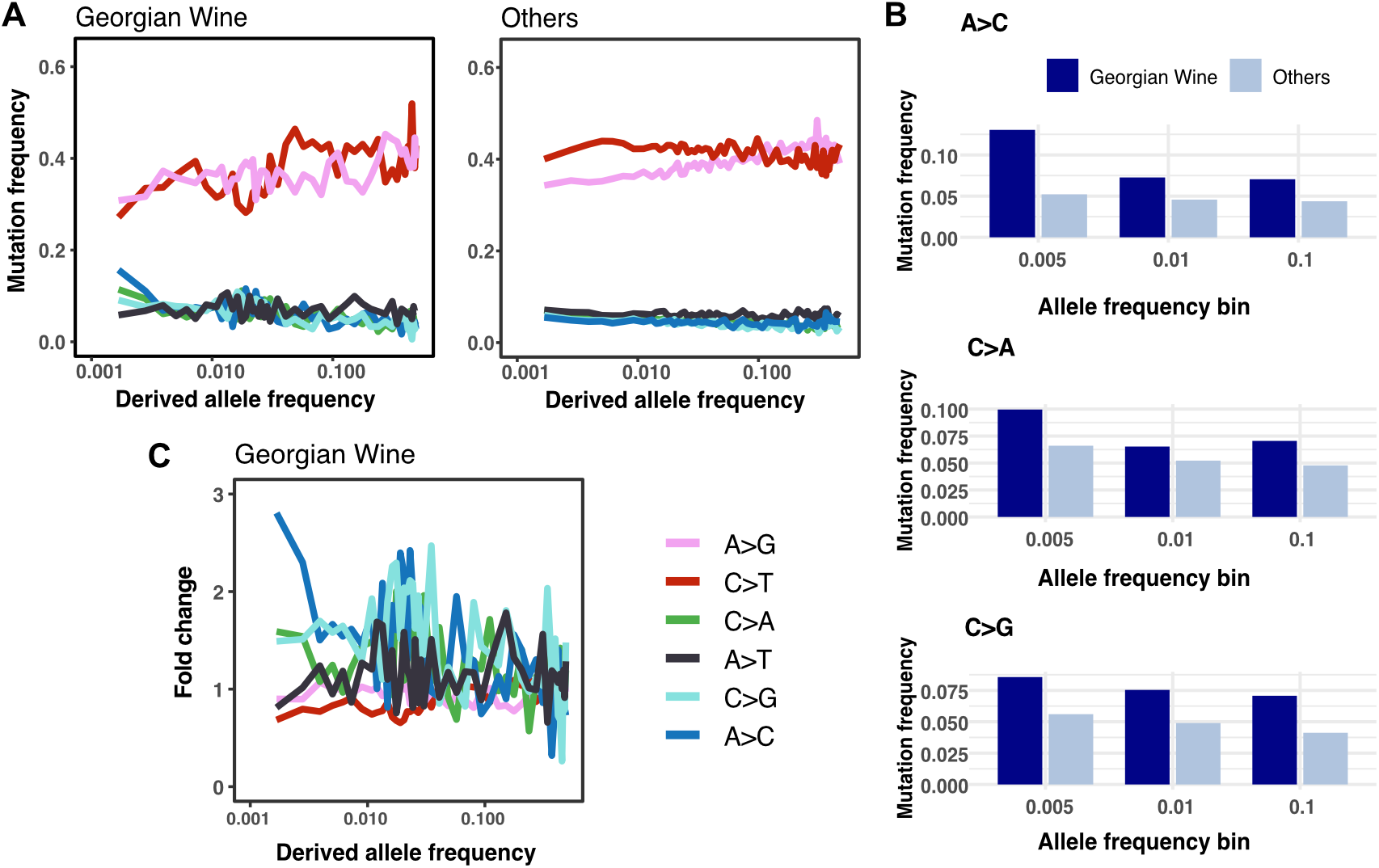
Georgian Wine population exhibits an outlier mutation spectrum. (A) Mutation frequency across allele frequency of Georgian Wine, and Others (combining alleles from all remaining populations except for African Beer, French dairy, and groups with extensive introgression from *S. paradoxus*, Alpechin, Mexican Agave, and French Guiana Human). (B) Georgian Wine population shows enrichment of transversion mutation types A>C, C>A, and C>G, among allele frequency bins. The enrichment is most prominent in the rarest allele frequency bin. (C) Fold change in allele frequency for Georgian Wine normalized to Others. A>C, C>A, and C>G showed 3-fold to 1.5-fold enrichment near the rarest allele frequency.

**Figure 4:**
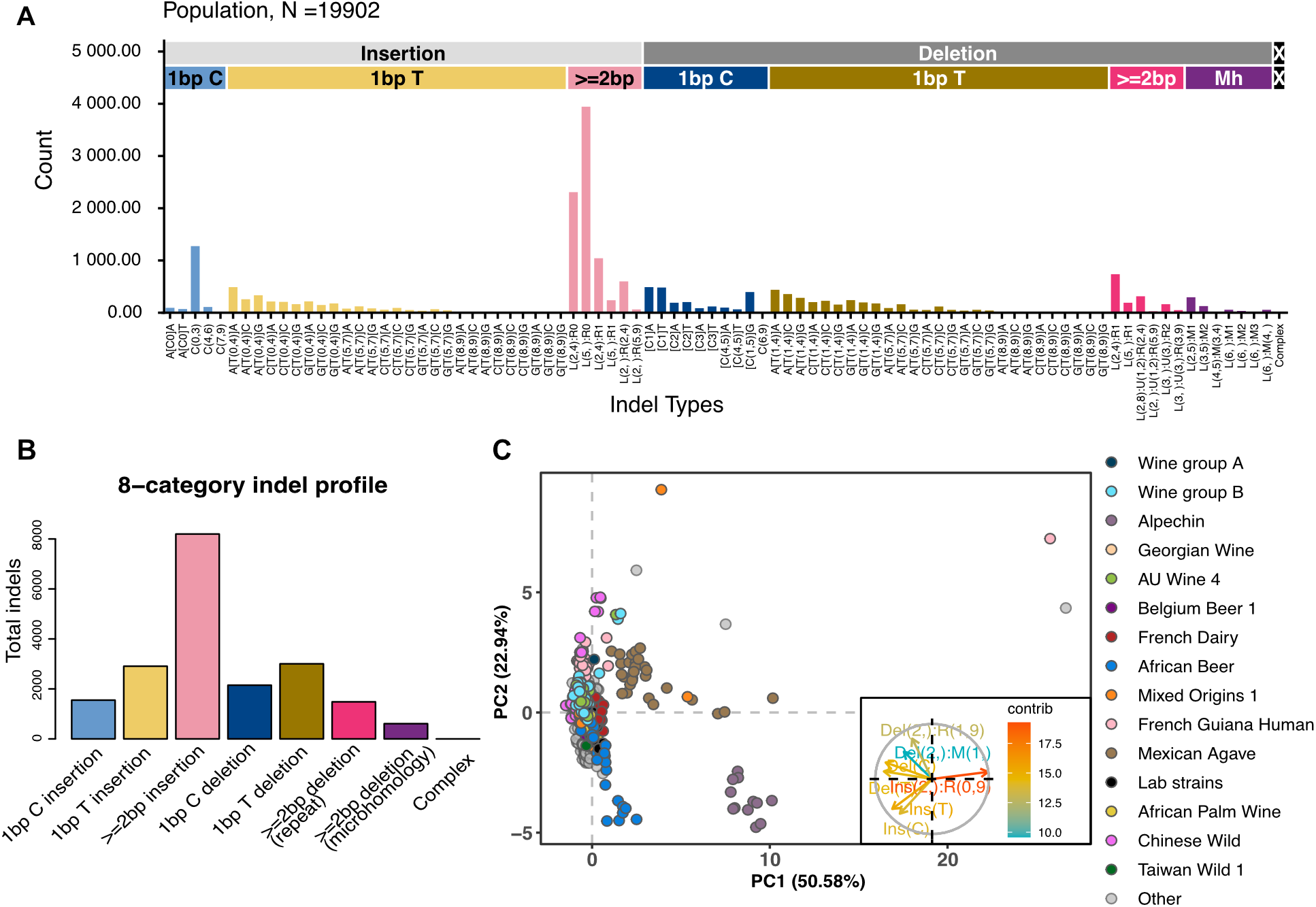
Indel classification and PCA highlight >=2 bp insertions: (A) Table of unique indel counts of *S. cerevisiae* classified via the 89 channels used in (Koh et al. 2025). 1 bp indels are split into C and T homopolymers due to the double-stranded nature and account for increased A/T slippage or C/G repeat instability. (B) Collapsed representation of all unique indel mutations into eight broad categories. (C) PCA of *S. cerevisiae* indel counts using the same eight categories.

We further analyzed the mutation-spectrum variation using rare genetic variants (Figure 2B). Consistent with the previous study, we found substantially less clustering of strains based on rare variants than based on common variants (Jiang et al. 2021). Several strains previously identified as having outlier mutation spectra also appeared as outliers in this analysis. For example, AAR and AEQ strains were discovered to exhibit elevated C>A mutations in rare polymorphisms and are also outliers along the correct axis in this analysis (Figure 2B). In addition, we observed outlier mutation spectra among several strains from the wine group that were not included in the previous analysis. These included XTRA_FPQ and XTRA_DMY from Wine group B and two AU Wine 4 strains, XTRA_GBV and XTRA_FEF, all of which were additional samples in the 3,034-genome dataset.

### Georgian Wine groups exhibited outlier mutation spectra among the 3034 *S. cerevisiae* genomes

The Georgian Wine stood out and displayed an outlier mutation spectrum along the PC1 axis (Figure 2A). The PCA loadings indicated that this group was enriched for C>A, C>G, and A>C mutation types. Compared with the SigProfiler results, C>A was the only enriched mutation type shared by SBS32 and the PCA loadings. One possible explanation is that COSMIC signatures were derived primarily from human cancers and may not capture additional mutational processes present in yeast populations. Nevertheless, the observed pattern may still reflect novel mutational processes operating in yeast populations.

To further investigate the evolutionary history of mutation-spectrum changes in Georgian Wine, we tracked the frequencies of different mutation types across allele-frequency bins, which provide a rough proxy for allele age. Rarer alleles are, on average, more likely to have arisen recently. We observed pronounced increases in A>C, C>A, and C>G mutations in Georgian Wine in the rarest allele-frequency bin (allele frequency <0.005, Figure 3A, 3B). Strikingly, we observed almost a 3-fold elevation of A>C for the rarest allele frequency, compared with ~1.5-fold elevation for C>A and C>G (Figure 3C). The 3-fold elevation is notable because, for a recent analysis with African Beer regarding its allele frequency change in the 1011 strains, a close to 2-fold elevation was observed for A>C at intermediate allele frequencies (Supplementary Figure 8). African Beer population has been the most prominent outlier in the mutational PCA analysis. Interestingly, the mutation elevation in Georgian Wine is pronounced among the rarest variants, but for A>C and C>G, there was a hump at intermediate frequencies, suggesting that additional mutator alleles or transient environmental change during the population’s history could have contributed to the intermediate mutation frequency signal.

### Indels from 3034 *S. cerevisiae* show non-uniform distribution across types and strains

We extended our mutation-spectrum analysis to indels in the 3,034 *S. cerevisiae* strains. Indels were polarized using the *S. paradoxus* CBS432 genome to infer ancestral states (Materials and Methods). To identify overall patterns and population-specific differences, we classified indels using IndelSigTools (Koh et al. 2025), a recently developed classification framework that has been applied to human genomes. We first assigned indels to 89 categories optimized by human indel types following Koh et al. (2025) and visualized their distribution across 1,793 non-closely related *S. cerevisiae* strains (Figure 4A). Because counts were sparse in many of these refined categories, we further collapsed them into eight broad classes: 1-bp C insertions, 1-bp T insertions, ≥2-bp insertions, 1-bp C deletions, 1-bp T deletions, ≥2-bp deletions, microhomology-associated deletions, and complex indels (Figure 4B). Among these classes, ≥2-bp insertions were the most abundant (Figures 4A, 4B). Within this class, non-repetitive insertions of 2-4 bp and ≥5 bp were the two most abundant categories (Figure 4A). Among ≥2-bp deletions, non-repetitive deletions of 2-4 bp were the most abundant, whereas among single-base insertions, C insertions associated with tracts of 0-3 repeat units were particularly common. We detected no complex indels in the dataset. Although indels were distributed across many channels, 1-bp insertions and deletions associated with longer repeat tracts were rare or absent.

We next performed PCA using indel counts collapsed into the eight broad categories because counts were sparse across the full set of 89 categories, and which is also human-centric. PC1 was dominated by ≥2-bp insertions, which contributed strongly to its positive direction (Figure 4C). The most prominent positive-PC1 outliers were strains from populations with substantial *S. paradoxus* introgression. In particular, Alpechin and Mexican Agave strains showed an excess of ≥2-bp insertions, while one French Guiana Human strain was also strongly displaced along this axis. These introgressed strains showed greater differentiation in their indel profiles than in their SNP spectra (Figures 2A and 4C). This stronger separation may reflect greater indel than SNP turnover since the divergence of *S. cerevisiae* and *S. paradoxus*, although it is possible that indels are subject to more errors during ancestral state polarization.

Among the remaining populations, most strains clustered near the origin. Several additional positive-PC1 outliers were observed, including two Mixed Origins 1 strains, two Wine group B strains, one AU Wine 4 strain, and three strains assigned to the Other group. These pronounced outliers may obscure subtler shifts in African Beer and Chinese Wild strains. Although PC2 was less readily interpretable, its loadings suggested possible enrichment of 1-bp T or C insertions in African Beer and of ≥2-bp deletions in Chinese Wild and several individual wine and Mixed Origins 1 outliers. The modest displacement of African Beer provides a candidate signal for investigating concordant mutational changes between SNVs and indels in that population.

## Discussion

In this study, we expanded the analyses of mutation-spectrum variation in natural *S. cerevisiae* isolates to a collection of 3,034 genomes, the largest population genomic dataset in *S. cerevisiae* to date. This collection includes novel groups not represented in the 1,011-genome dataset, in addition to denser sampling and finer population assignments for previously represented groups. The overall pattern of mutation-spectrum variation was largely consistent with that observed in the 1,011-genome collection, indicating that the major population-level differences are robust to expanded sampling. At the same time, the newly included strains and improved population resolution revealed distinct mutational patterns within the wine superclade and identified Georgian Wine as an additional population-level outlier.

The finer division of the wine superclade revealed a novel mutation signature involving SBS32, which is not present in the 1,011-genome collection (Jiang et al. 2025). Most Wine group A strains exhibited SBS32 in place of SBS102, whereas most Wine group B strains retained SBS102 and lacked SBS32. This variation indicates that the wine superclade does not share a single mutation spectrum. Wine group A contains Georgian Wine along with AU Wine 1-3, and West Europe Wine contained such a novel signature.

Interestingly, when performing mutational PCA analysis on the extended dataset, only Georgian Wine stood out as an outlier among the wine superclade, with C>A, C>G, and A>C enrichment. Upon further scrutiny with allele frequencies, we discovered that Georgian Wine is enriched in all three mutation types, especially among the rarest allele frequencies. This suggests that these mutation rate increases in Georgian Wine are most likely to have occurred in the recent history. Substantial mutation frequency increases were observed, with an almost threefold elevation of A>C mutations. The threefold elevation of A>C enrichment is even higher than the difference observed in the most striking outlier, African Beer, where an approximately twofold elevation was most apparent at intermediate allele frequencies. Plausible causes of this pattern could be novel mutator alleles that arose in the Georgian Wine background, or some recent environmental shift that affects mutation spectrum changes in the Georgian Wine.

Our indel analysis suggests that population-level mutation-spectrum variation in *S. cerevisiae* extends beyond single-nucleotide substitutions. Insertions of ≥2 bp were the most abundant class of unique indel variants, and the indel PCA identified African Beer as an outlier. This partial correspondence with the SNP analysis raises the possibility that lineage-specific differences in DNA replication or repair affect multiple mutation classes, although the underlying mechanisms remain to be tested. The *S. paradoxus*-introgressed Alpechin, Mexican Agave, and French Guiana Human populations showed a different pattern, characterized by an excess of insertions of ≥2 bp and stronger separation than in the SNP analysis. Because indel detection and ancestral-state polarization are particularly sensitive to repeat context, alignment, and reference divergence, especially in introgressed regions, this signal should be interpreted cautiously. It is likely the *S. paradoxus*-introgressed strains simply inherit more ancestral indel types with are enriched for ≥2 bp insertions. Nevertheless, these patterns serve as complementary information for strains in which direct measurements of de novo indels could test for underlying differences in indel formation and repair (Harfe and Jinks-Robertson 2000; Gragg et al. 2002). Together, these findings show that expanded sampling preserves the major patterns observed previously while revealing additional population-associated variation in both substitution and indel spectra.

## Acknowledgements

This work was supported by institutional research funds provided by the College of Liberal Arts and Sciences and Biodesign Institute of Arizona State University to PJ. No external funding was received for this project.

## Materials and Methods

### Variant filtering and mutation count

We performed our analysis with the filtered VCF file from 3034 natural isolates of *S. cerevisiae* (Loegler et al., 2024). Similar filtering and analysis steps follow previous work (Jiang et al., 2021), with slight modifications. Only biallelic SNP sites with a derived allele frequency of <0.5 were used, by excluding sites with >20% missing genotype data. Bcftools (version 1.14) and bedtools (version 2.30.0) were used to remove repeat-masked regions and retain SNPs located within uniquely mapped genomic regions (‘mregions_100_annot_2011.bed’ from Jubin et al., 2014). After excluding closely related strains (pairwise genetic distance less than 8000), there were 1793 remaining strains that were further subset to 858 after taking a maximum of 30 from each population. Another subset excludes strains that show extensive introgression, where 785 strains remain. Polymorphisms were polarized with *S. paradoxus* (Jiang et al., 2021). Both six-type single-nucleotide and 96-type trinucleotide (single-nucleotide mutations with immediate flanking 5’ and 3’ bases) mutations are included.

For rare variants, we raised the allele count cutoff from 4 to 12 due to increased sample size (from 1011 to 3034). We excluded singletons to minimize sequencing error impact.

### SigProfiler analysis

For each sample, mutation counts were aggregated across the 96 trinucleotide contexts to generate a 96×785 mutational count matrix, excluding introgressed strains. We then used SigProfilerExtractor v1.2.6 and SigProfilerMatrixGenerator v1.3.6 with default parameters (Islam et al., 2022) to decompose the observed mutational profiles into contributions from known COSMIC single base substitution (SBS) signatures.

### Mutational PCA analysis

Mutation counts were used for principal component analysis via prcomp for the mutational spectra of SNPs. The procedure closely followed previous work (Jiang et al., 2021), with the modification of the “is_scale=TRUE” flag for normalization.

### Georgian Wine mutation frequency

Mutation frequency from each type is binned by derived allele frequencies and analyzed separately for a population under study following (Jiang et al. 2025). The baseline “Others” group consisted of all strains excluding the Georgian Wine, African Beer, French Dairy, Alpechin, Mexican Agave, and French Guiana Human.

### Indel classification

The Indel VCF data was obtained from the 3034 genomes (Loegler et al. 2024). To infer the indels from an ancestral state, we replaced the reference genome with *S. paradoxus* CBS432 using bcftools + liftover BCFtools v1.22 / htslib v1.24 (Genovese et al. 2024).

Downstream indel mutation classification and counting is performed with indelsigtools package (Koh et al. 2025) using unique indels in the population. Indel mutation counts were initially classified into 476 or 89 categories using the established pipeline, and further grouped into eight broad categories: 1 bp T insertions, 1 bp T deletions, 1 bp C insertions, 1 bp C deletions, ≥2 bp insertions, ≥2 bp deletions, ≥2 bp deletions with microhomology, and complex. C and T-based classifications were used to account for the reverse complement.

### Indel PCA mutation spectra analysis

Individual indel mutation counts from eight broad categories were input into PCA. The complex indel category contained no mutations and was therefore excluded prior to PCA due to lack of variance. We performed indel PCA on the 858 strains made a cutoff of 350 minimum counts to remove any distortions from rare outliers. PCA via prcomp was then performed with default parameters and is_scale set to true.

