## Supplementary Figure 1-8 for "Mutation spectra across 3,034 *Saccharomyces cerevisiae* genomes distinguish the Georgian Wine population"

### Supplementary Figures

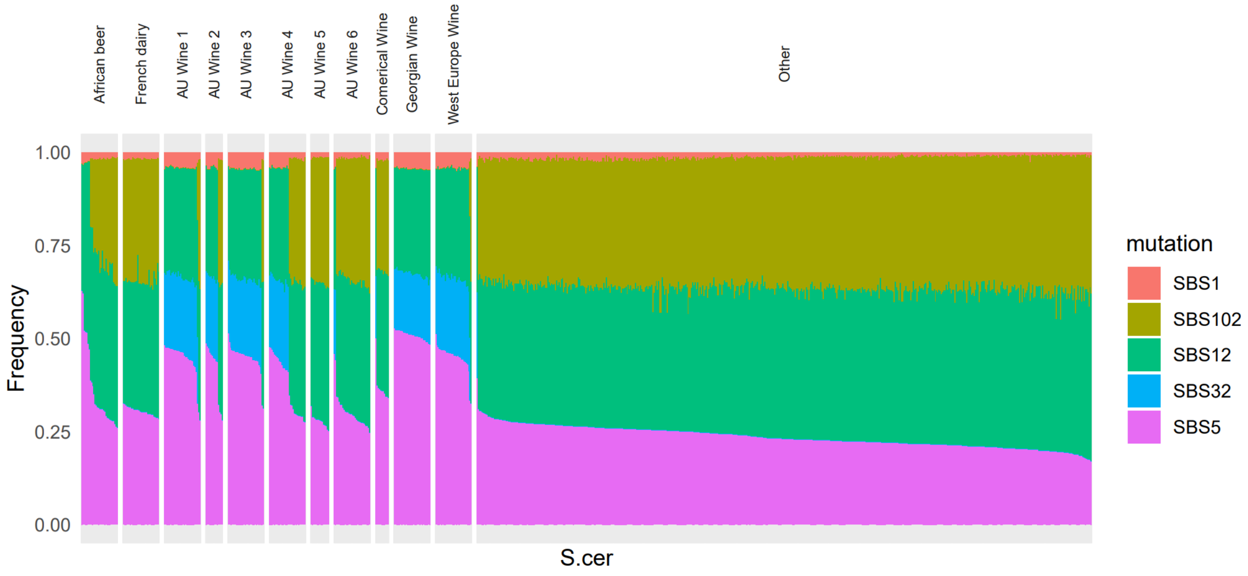


**Supplementary Figure 1. Wine Group Signature Decomposition.** Signature decomposition via SigProfiler. With the appearance of new signatures in the wine superclade, we separated the clades to determine which followed each pattern. This is how the wine groupings were determined. Wine group A (AU Wine 1-3, Georgian Wine, and West Europe Wine) exhibits differing signatures than Wine group B (AU Wine 5-6, and Commercial Wine) while AU Wine 4 has a relatively even split between signature profiles.

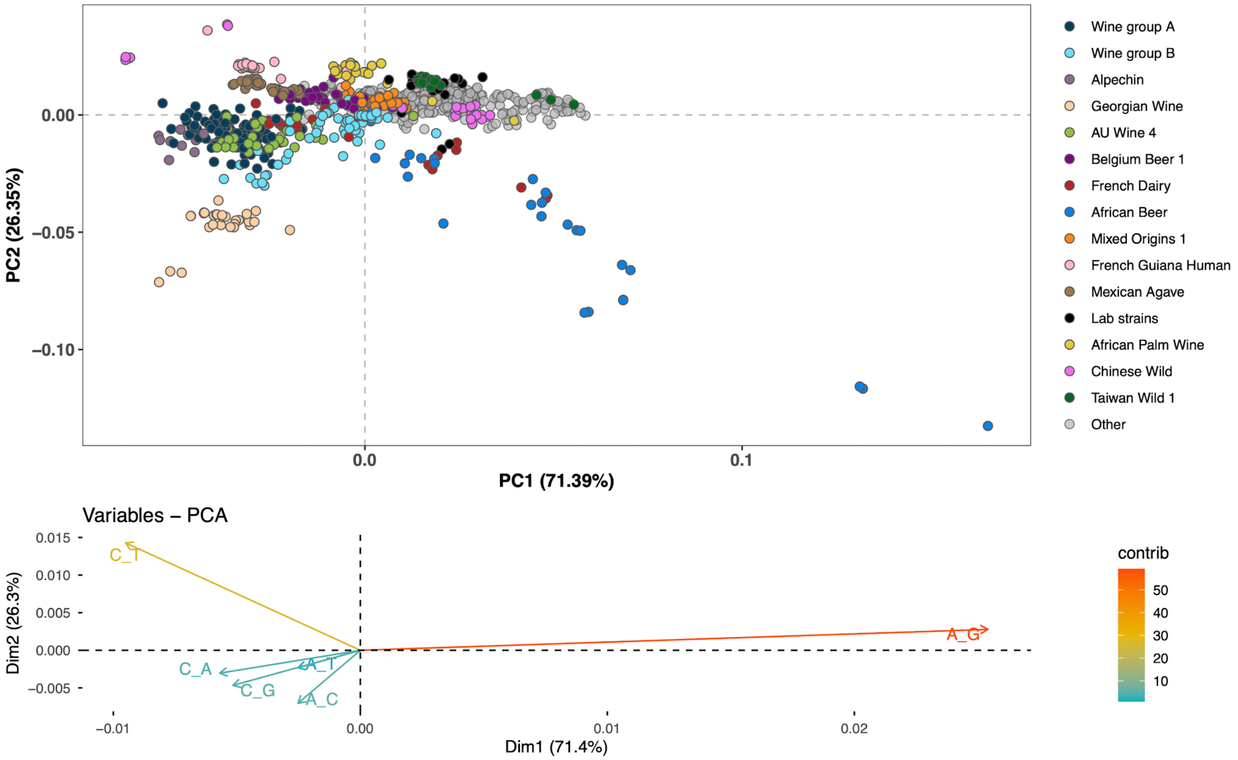


**Supplementary Figure 2.** ***S. cerevisiae*** **Mutation spectra without normalization show heavy A>G skew from African Beer.** *S. cerevisiae* mutation spectra before normalization. Heavy skewing from African beer strains drives the PC1 axis. These strains are enriched heavily with A>G substitutions.

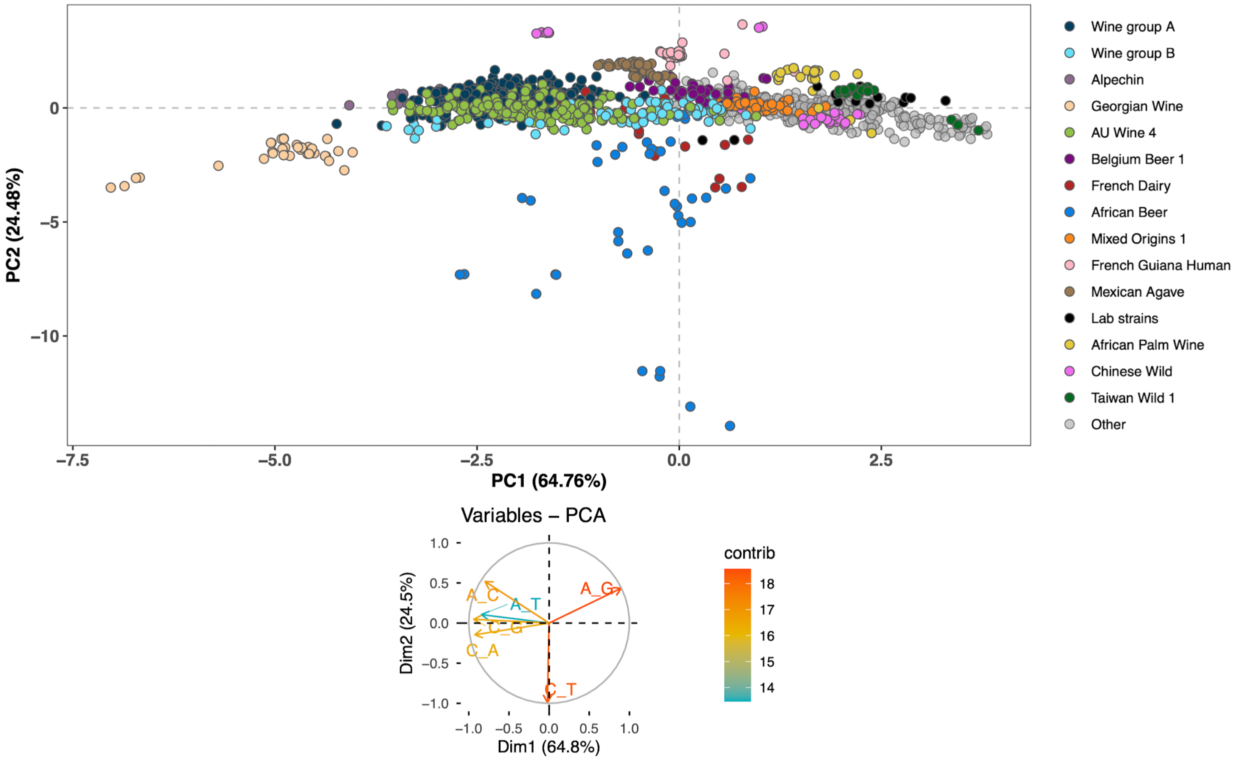


**Supplementary Figure 3. *S. cerevisiae* Mutation spectra of the 1793 non-closely related strains without subsampling.**

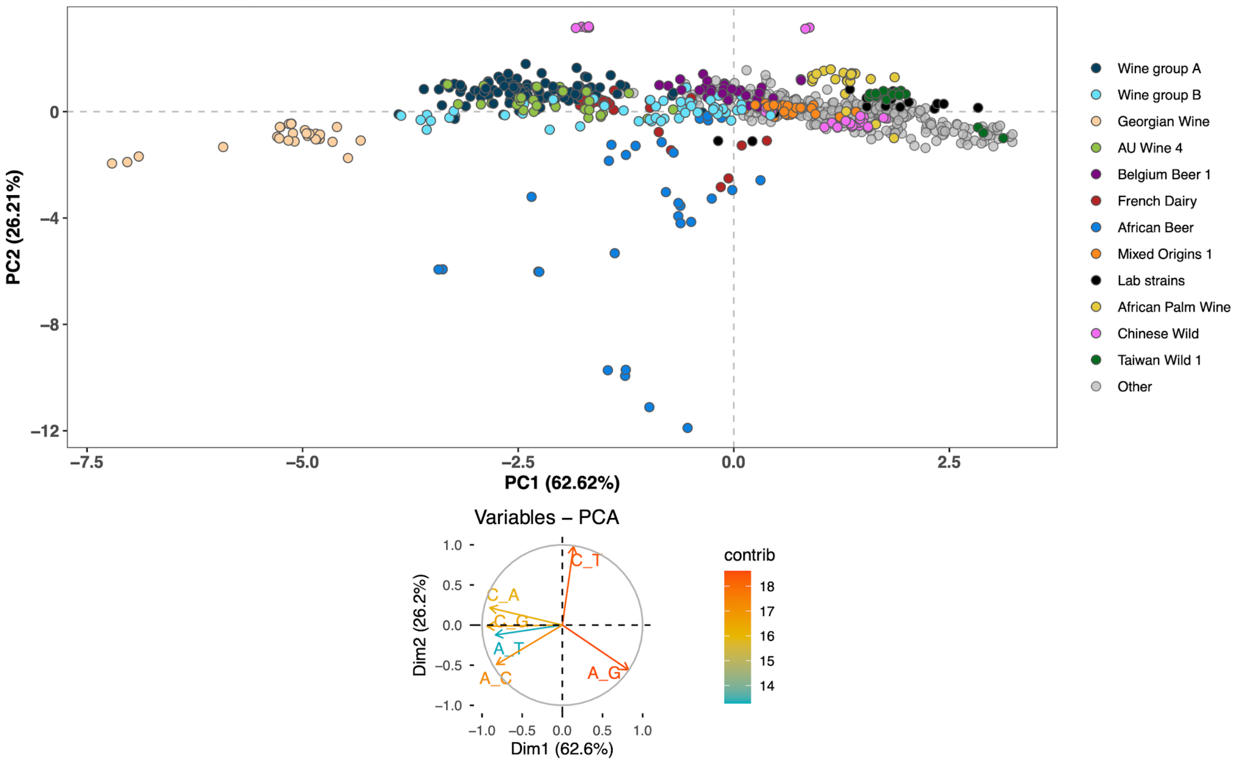

**Supplementary Figure 4.** ***S. cerevisiae*** **Mutation spectra without introgression:** Removes strains that exhibit significant introgression from the PCA (Alpechin, Mexican Agave, French Guiana Human). Patterns remain consistent, suggesting minimal interference caused by introgression.

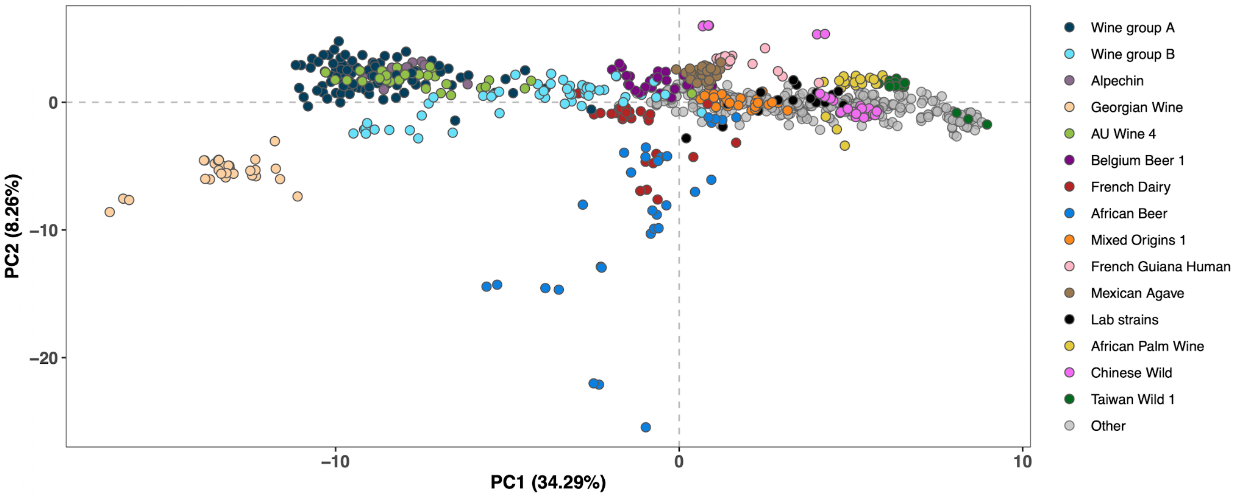


**Supplementary Figure 5.** ***S. cerevisiae*** **triplet mutation spectra:** Follows same default parameters as 2A, normalized 785 strain subset including introgressed strains. Same patterns can be seen, though Georgian Wine is even more of an outlier as it is further down the PC2 axis.

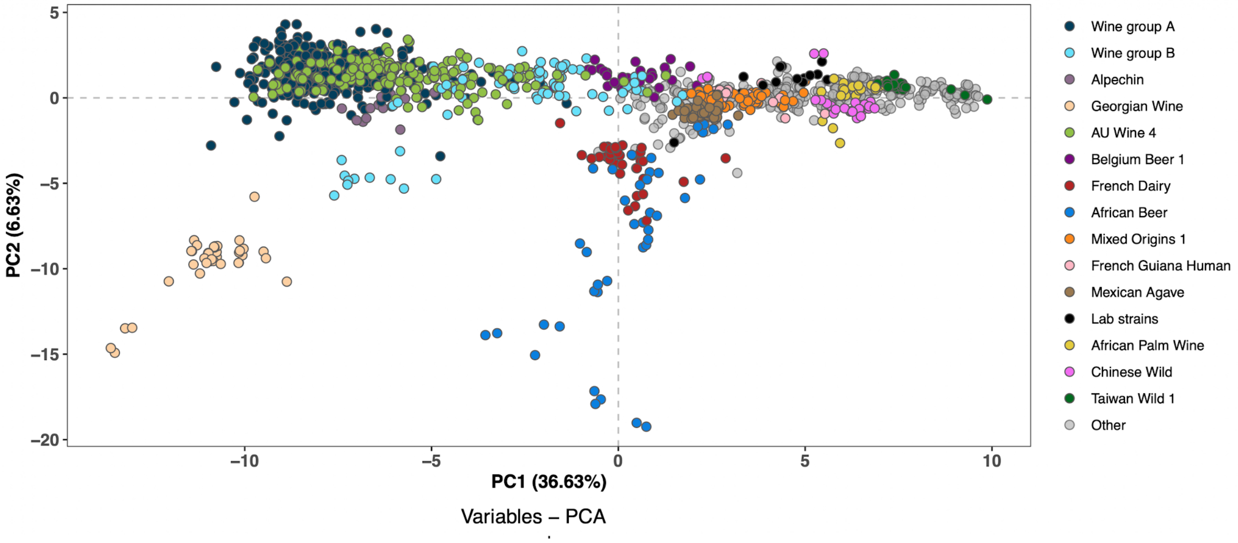


**Supplementary Figure 6. Triplet PCA among all 1793 non-closely related strains.**

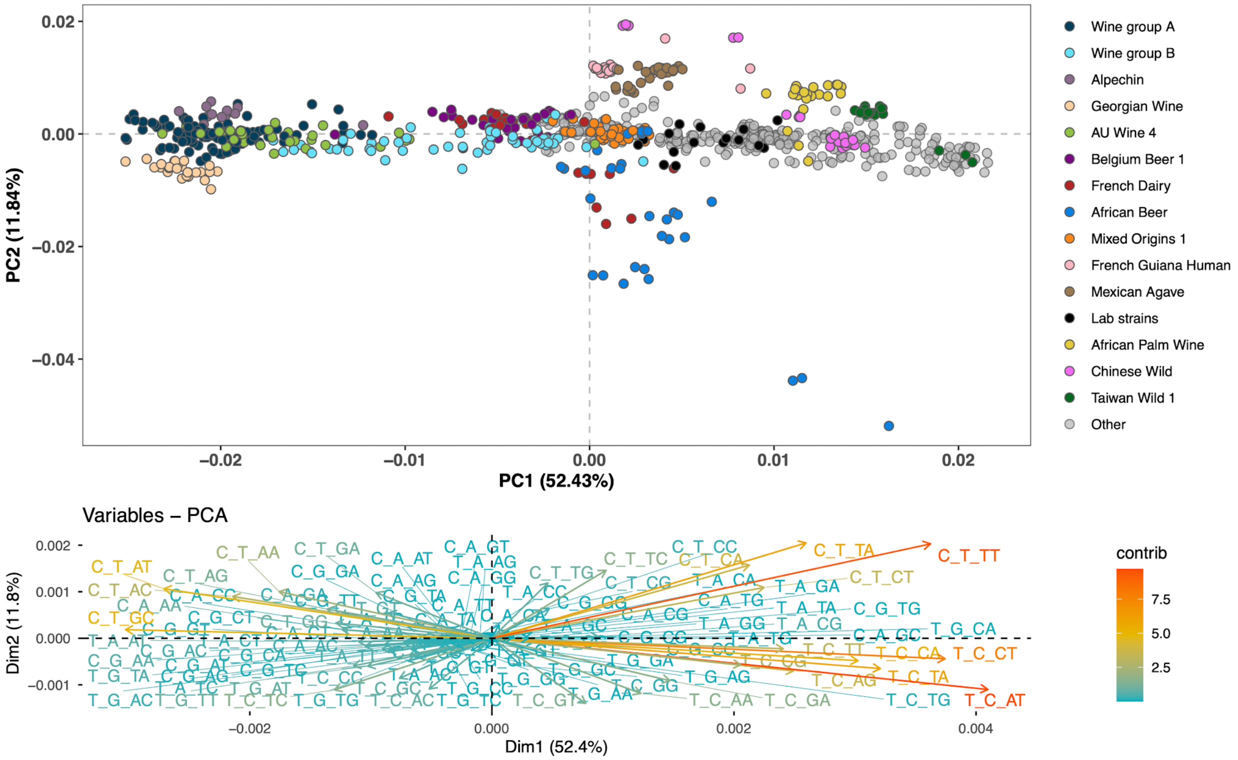


**Supplementary Figure 7. *S. cerevisiae* triplet mutation spectra without normalization.**

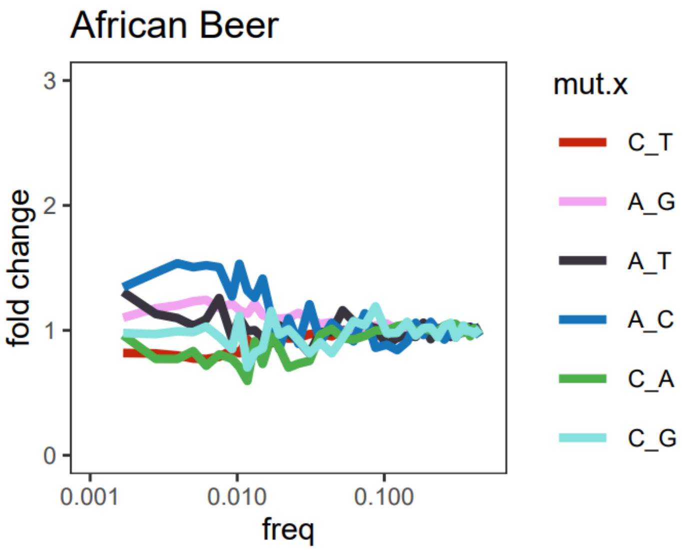


**Supplementary Figure 8. African Beer fold change in the 3034 dataset.**
